# The telegraph process is not a subordinator

**DOI:** 10.1101/2023.01.17.524309

**Authors:** Gennady Gorin, Lior Pachter

**Affiliations:** Division of Chemistry and Chemical Engineering, California Institute of Technology, Pasadena, CA, 91125; Division of Biology and Biological Engineering, California Institute of Technology, Pasadena, CA, 91125; Department of Computing and Mathematical Sciences, California Institute of Technology, Pasadena, CA, 91125

## Abstract

Investigations of transcriptional models by Amrhein et al. outline a strategy for connecting steady-state distributions to process dynamics. We clarify its limitations: the strategy holds for a very narrow class of processes, which excludes an example given by the authors.

## 1 BACKGROUND

A preprint by Amrhein et al. (1), adapted into Ch. 4 of the dissertation (2), describes the class of transcription and degradation processes:

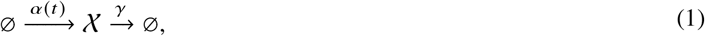

where 𝒳 is an RNA transcript, *α*(*t*) is its transcription rate, and *γ* is its degradation rate. *α*(*t*) may be stochastic, deterministic, or constant. The distribution *P* of the discrete counts of 𝒳 is given by a Poisson mixture, such that

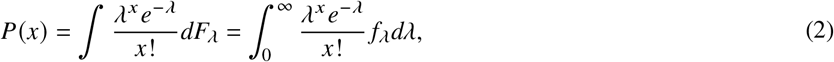

where *λ* is a mixing parameter that has a probability distribution function *f*_*λ*_. The time-dependent distribution of *λ* can be obtained by solving the underlying stochastic differential equation:

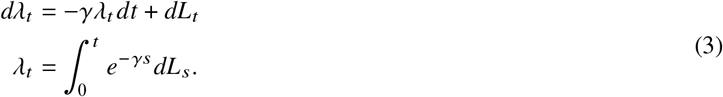

This follows from the Poisson representation (3, 4), which has been applied to analogous problems (5, 6). Informally, *dL*_*t*_ is the instantaneous contribution from the transcription rate process, e.g., *α*(*t*) *dt* if *α* is deterministic.

Amrhein et al. note that if *L*_*t*_ is a *subordinator*, the stationary law of *f*_*λ*_ can be obtained by straightforward manipulations (7) and that furthermore, this stationary law is *self-decomposable*. Conversely, every self-decomposable law can be represented as the stationary distribution of a process driven by some subordinator.

In this context, a process is a subordinator if it is Lévy and increasing. The Lévy property requires stationary and independent increments (7). A self-decomposable law is one that has the property *G*(*z*) = *G*(*cz*) *G*_*c*_(*z*) for all *c* ∈ (0, 1), where *G*(*z*) is the law’s characteristic function and *G*_*c*_ is another characteristic function. If these criteria are met, then

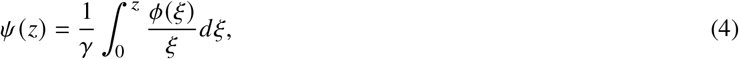

where *ψ*(*z*) is the log-characteristic function of the stationary distribution of *λ*, and *ϕ* is the log-characteristic function of the subordinator *L*_*t*_ at *t* = 1.

Finally, Amrhein et al. assert that the *telegraph model* can serve as such a subordinator (e.g., Fig. 2 and p. 6 of (1)). The telegraph model describes transitions between two states (“on” and “off”), such that the transcription rate in the on state is *k*_*tx*_ (8). The steady-state distribution of the corresponding process is Poisson-Beta, i.e., the underlying continuous process has a Beta stationary law (9). The notation suggests that the process governing the Beta-distributed *λ* can be cast in the form of Equation 3, i.e., a single stochastic differential equation driven by a subordinator. Specifically, Amrhein et al. define the *integrated telegraph process* 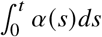, such that *α*(*s*) = *k*_*tx*_ if the gene switch is in the “on” state and 0 otherwise, and propose that it constitutes a subordinator. However, Amrhein et al. do not proceed to use the approach in Equation 4 to obtain the stationary distribution, opting to follow a different derivation (5).

## 2 RESULTS

The Amrhein et al. manuscript frames the connection between stochastic differential equations and chemical master equations as its key result, uses the same notation for all described processes, and explicitly asserts that the telegraph process can be represented in terms of a subordinator. It can therefore potentially be misleading, in that it suggests that the procedure in Equation 4 applies to the telegraph process. This implication is incorrect. The procedure is legitimate for compound Poisson (Sec. 4.4.2 and Sec. 4.3.1 of (2)) subordinators, among others (Supp. Sec. 5.3 of (10)). However, the relevant telegraph-derived process (realization shown in the left panel of Fig. 2 of (1)) is *not* a subordinator, and cannot be represented in the form of Equation 3. We present three arguments for why this is the case.

### Distribution class

The steady state of the telegraph model is Beta-Poisson. Its mixing density is Beta (9). All subordinator-driven Ornstein-Uhlenbeck processes induce self-decomposable stationary laws (11). All self-decomposable laws are unimodal (12). Unimodal mixing distributions yield unimodal Poisson mixtures (13). Since the Beta-Poisson distribution may be bimodal (Figure 1a), the underlying bimodal Beta law is not self-decomposable, implying the integrated telegraph process is not a subordinator.

**Figure 1:**
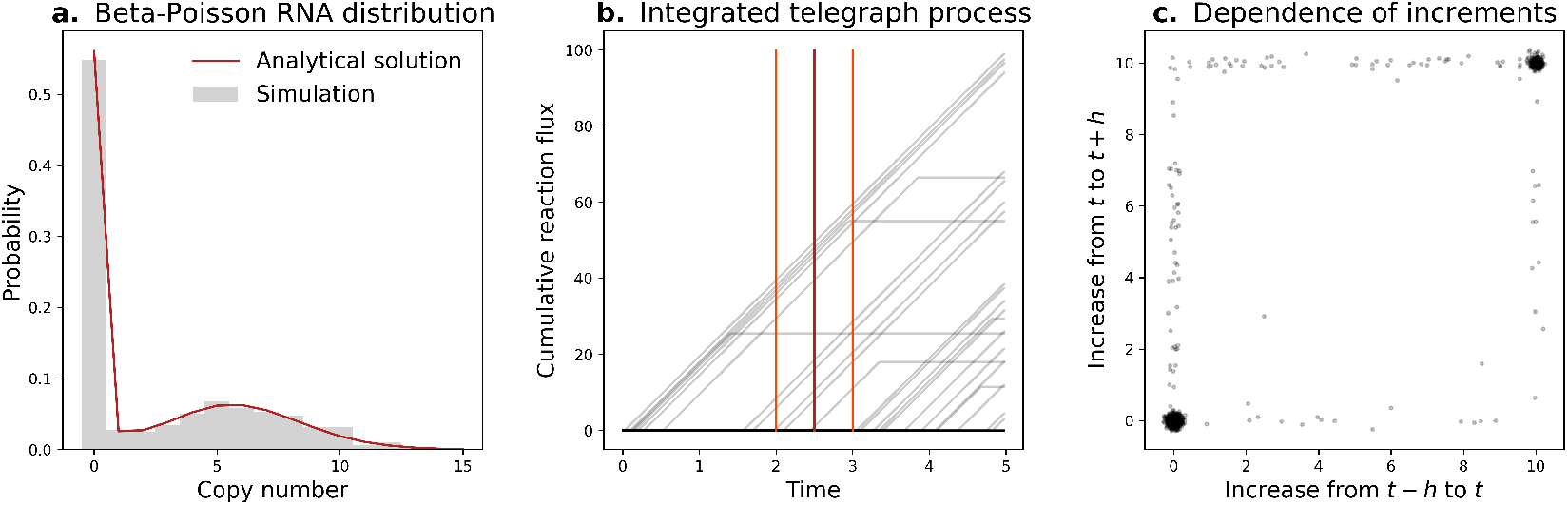
The telegraph process is not a subordinator. **a**. The stationary distribution is bimodal, implying the mixing distribution is not self-decomposable (histogram: 1,000 simulated realizations; red line: analytical solution (15, 16)). **b**. The trajectory shapes (gray lines) disagree with Lévy criteria (fifty realizations shown; dark red: reference time *t*; orange-red: time points *t* + *h* and *t* − *h*). **c**. Disjoint increments are non-independent (points: 1,000 simulated realizations).

### Admissible trajectory shapes

The integrated telegraph process is continuous and almost everywhere differentiable (Figure 1b). The only continuous Lévy processes are the Brownian motions with drift (14). The only continuous and differentiable Lévy processes have the structure *X*_*t*_ = *kt*, implying the integrated telegraph process is not a subordinator and the premise does not hold.

### Increment conditions

A subordinator has independent increments (14). The integrated telegraph process fails to meet this criterion: the evolution of the process from time *t* to *t* + *h* is strongly dependent on its evolution from *t* − *h* to *t*. In the most striking case, if the switching rates are much lower than *h*^−1^, the two segments become highly correlated (Figure 1c). Therefore, this process is not a subordinator and the premise does not hold.

## Conclusion

The integrated telegraph process happens to *converge to* the trivial *kt* subordinator in the constitutive limit and the compound Poisson subordinator in the bursty limit. However, generally, representing driving by stochastic processes necessitates explicitly coupling these processes to the chemical master equation, and requires considerable analytical effort (10).

## 3 METHODS

To generate synthetic data for Figure 1, we simulated a system with *k*_*on*_ = 0.15, *k*_*off*_ = 0.1, *k*_*tx*_ = 20, and *γ* = 3.14 using Gillespie’s stochastic simulation algorithm (17), as previously implemented for (18). We performed 1,000 simulations, run until *t* = 5, with the system state stored at 200 uniformly spaced time points (Δ*t* = 0.025).

For the analytical solution in Figure 1a, we used the results from Huang et al. (15), setting the feedback term to zero. This implementation was previously used for (16, 19). To obtain the “subordinator” functions for Figure 1b, we computed the integral of the observed transcription rates, *Y*_*t*_. This quantity is the cumulative reaction flux of the transcription reaction up to a given time. The panel shows the reference time *t* = 2.5, and the increment bounds *t* + *h* and *t* − *h*, with *h* = 0.5. In Figure 1c, we plot the value of *Y*_*t*+*h*_ − *Y*_*t*_ against the value of *Y*_*t*_ − *Y*_*t*−*h*_. The visualization includes Gaussian jitter with *σ* = 0.1. For a process with independent increments, the distribution of these quantities must be independent.

## 4 CODE AVAILABILITY

The Python notebook used to generate Figure 1 is available at https://github.com/pachterlab/GP_2023.

## 5 ACKNOWLEDGMENTS

G.G. and L.P. were partially funded by NIH U19MH114830. The Huang et al. Poisson-Beta solution was implemented in Python by Dr. John J. Vastola, who contributed valuable insights in our broader investigation of continuous and discrete master equations.

## REFERENCES

1. Amrhein, L., K. Harsha, and C. Fuchs, 2019. A mechanistic model for the negative binomial distribution of single-cell mRNA counts. Preprint, bioRxiv: 657619. http://biorxiv.org/lookup/doi/10.1101/657619.

2. Amrhein, L., 2021. Stochastic Modeling of Heterogeneous Low-Input Gene Expression: Linking Single-Cell Probability Distributions to Transcription Mechanisms. PhD Dissertation, Technische Universitat Munchen, Munich.

3. Gardiner, C. W., and S. Chaturvedi, 1977. The poisson representation. I. A new technique for chemical master equations. Journal of Statistical Physics 17:429–468. http://link.springer.com/10.1007/BF01014349.

4. Gardiner, C., 2004. Handbook of Stochastic Methods for Physics, Chemistry, and the Natural Sciences. Springer, third edition.

5. Smiley, M. W., and S. R. Proulx, 2010. Gene expression dynamics in randomly varying environments. Journal of Mathematical Biology 61:231–251. http://link.springer.com/10.1007/s00285-009-0298-z.

6. Iyer-Biswas, S., and C. Jayaprakash, 2014. Mixed Poisson distributions in exact solutions of stochastic auto-regulation models. Physical Review E 90:052712. http://arxiv.org/abs/1110.2804, 1110.2804.

7. Barndorff-Nielsen, O. E., and N. Shephard, 2001. Non-Gaussian Ornstein-Uhlenbeck-based models and some of their uses in Financial economics. Journal of the Royal Statistical Society: Series B 63:167–241. https://rss.onlinelibrary.wiley.com/doi/10.1111/1467-9868.00282.

8. Peccoud, J., and B. Ycard, 1995. Markovian Modeling of Gene Product Synthesis. Theoretical Population Biology 48:222–234.

9. Stinchcombe, A. R., C. S. Peskin, and D. Tranchina, 2012. Population density approach for discrete mRNA distributions in generalized switching models for stochastic gene expression. Physical Review E 85:061919. https://link.aps.org/doi/10.1103/PhysRevE.85.061919.

10. Gorin, G., J. J. Vastola, M. Fang, and L. Pachter, 2022. Interpretable and tractable models of transcriptional noise for the rational design of single-molecule quantification experiments. Nature Communications 13:7620. https://www.nature.com/articles/s41467-022-34857-7.

11. Barndorff-Nielsen, O. E., and S. Thorbjørnsen, 2002. Self-Decomposability and Lévy Processes in Free Probability. Bernoulli 8:323–366. http://www.jstor.org/stable/3318705.

12. Yamazato, M., 1978. Unimodality of Infinitely Divisible Distribution Functions of Class L. The Annals of Probability 6:523–531. http://www.jstor.org/stable/2243119, publisher: Institute of Mathematical Statistics.

13. Karlis, D., and E. Xekalaki, 2005. Mixed Poisson Distributions. International Statistical Review / Revue Internationale de Statistique 73:35–58. http://www.jstor.org/stable/25472639.

14. Barndorff-Nielsen, O. E., S. I. Resnick, and T. Mikosch, editors, 2001. Lévy Processes. Birkhäuser Boston, Boston, MA. http://link.springer.com/10.1007/978-1-4612-0197-7.

15. Huang, L., Z. Yuan, P. Liu, and T. Zhou, 2014. Feedback-induced counterintuitive correlations of gene expression noise with bursting kinetics. Physical Review E 90:052702. https://link.aps.org/doi/10.1103/PhysRevE.90.052702.

16. Vastola, J. J., G. Gorin, L. Pachter, and W. R. Holmes, 2021. Analytic solution of chemical master equations involving gene switching. I: Representation theory and diagrammatic approach to exact solution. Preprint, 2103.10992. http://arxiv.org/abs/2103.10992, 2103.10992.

17. Gillespie, D. T., 1977. Exact stochastic simulation of coupled chemical reactions. The Journal of Physical Chemistry 81:2340–2361. https://pubs.acs.org/doi/abs/10.1021/j100540a008.

18. Gorin, G., S. Yoshida, and L. Pachter, 2022. Transient and delay chemical master equations. Preprint, bioRxiv: 2022.10.17.512599. http://biorxiv.org/lookup/doi/10.1101/2022.10.17.512599.

19. Vastola, J. J., 2021. In search of a coherent theoretical framework for stochastic gene regulation. Ph.D. thesis, Vanderbilt. https://ir.vanderbilt.edu/handle/1803/16646.

